# Automated and Manual Quantification of Tumour Cellularity in Digital Slides for Tumour Burden Assessment

**DOI:** 10.1101/571190

**Authors:** Shazia Akbar, Mohammad Peikari, Sherine Salama, Azadeh Y. Panah, Sharon Nofech-Mozes, Anne L. Martel

## Abstract

**Aims:** The residual cancer burden index is an important quantitative measure used for assessing treatment response following neoadjuvant therapy for breast cancer. It has shown to be predictive of overall survival and is composed of two key metrics: qualitative assessment of lymph nodes and the percentage of invasive or in-situ tumour cellularity (TC) in the tumour bed (TB). Currently, TC is assessed through eye-balling of routine histopathology slides estimating the proportion of tumour cells within the TB. With the advances in production of digitized slides and increasing availability of slide scanners in pathology laboratories, there is potential to measure TC using automated algorithms with greater precision and accuracy.

**Methods:** We describe two methods for automated TC scoring: 1) a traditional approach to image analysis development whereby we mimic the pathologists’ workflow, and 2) a recent development in artificial intelligence in which features are learned automatically in deep neural networks using image data alone.

**Results:** We show strong agreements between automated and manual analysis of digital slides. Agreements between our trained deep neural networks and experts in this study (0.82) approach the inter-rater agreements between pathologists (0.89). We also reveal properties that are captured when we apply deep neural network to whole slide images, and discuss the potential of using such visualisations to improve upon TC assessment in the future.

**Conclusions:** TC scoring can be successfully automated by leveraging recent advancements in artificial intelligence, thereby alleviating the burden of manual analysis.

## 1 Introduction

Neoadjuvant systemic therapy (NAT) for breast cancer (BC) is used to treat locally advanced and operable BC to allow breast-conserving surgery [1]. NAT provides an opportunity to monitor clinical, radiological and ultimately with pathologic response. Pathologic complete response (pCR) to NAT has been shown to predict survival [2] and local recurrence [3]. As such, accurate assessment of pathologic response to NAT provides important prognostic information.

Standardized protocol to assess the extent of response with objective measures across multiple institutions is essential. Symmans et al. [4] proposed a method for quantifying residual disease by calculating the residual cancer burden index (RCB). RCB is recognized as long-term prognostic utility [4] and has shown to be more predictive of overall survival compared to other measurements [5]. The RCB index accounts for two key metrics: qualitative assessment of residual disease in the breast (via tumour cellularity (TC) in the tumour bed (TB) and proportion of in situ component) and assessment of lymph nodes). While RCB calculator produces a continuous score, scores are further categorized in four RCB classes from pCR (RCB-0) to extensive residual disease (RCB-III) that are easily reproducible [6]. Accurate quantification of TC is laborious, time consuming task that most practicing pathologists are not trained to perform. Yet, TC is crucial for computing the RCB index.

Currently TC is estimated by manually estimation (“eyeballing”) the TB area at multiple microscopic fields through several slides that represent the largest linear dimension, and comparing the involved area with graphic standard skatches [7]. Such illustrations although helpful are semi-quantitative, subjective measurements. A single case-level score is then obtained by taking an average of TC scores from different fields and rounded to the nearest 10th percent. However, these scores can also be reformulated on a continuous scale. As manual analysis is limited to predefined discrete values, we are yet to discover the potential benefits of continuous TC scores for prognosis. In theory such measurements can be more precise, however, acquiring them manually is infeasible given there is a greater chance of error and reproducibility is not possible.

With latest technological advancement in digital pathology, including tissue scanners capable of scanning whole slides at high resolutions, there is potential to leverage image analysis techniques to gain more accurate metrics than is achievable by the human eye, and reduce pathology workload by eliminating time-consuming tasks. TB region can be captured digitally on scanned slides, using annotation tools (Figure 1). Towards an ultimate goal to automate RCB assessment we explored methods to evaluate TC as the first step. In this paper, we report the use of automation to compute TC scores using two different image analysis approaches (Figure 2):

1. An image analysis pipeline which encompasses hand-engineered features designed to mimic the pathologist’s eye.
2. A deep learning approach which learns features directly from raw image data of digital slides.

**Figure 1:**
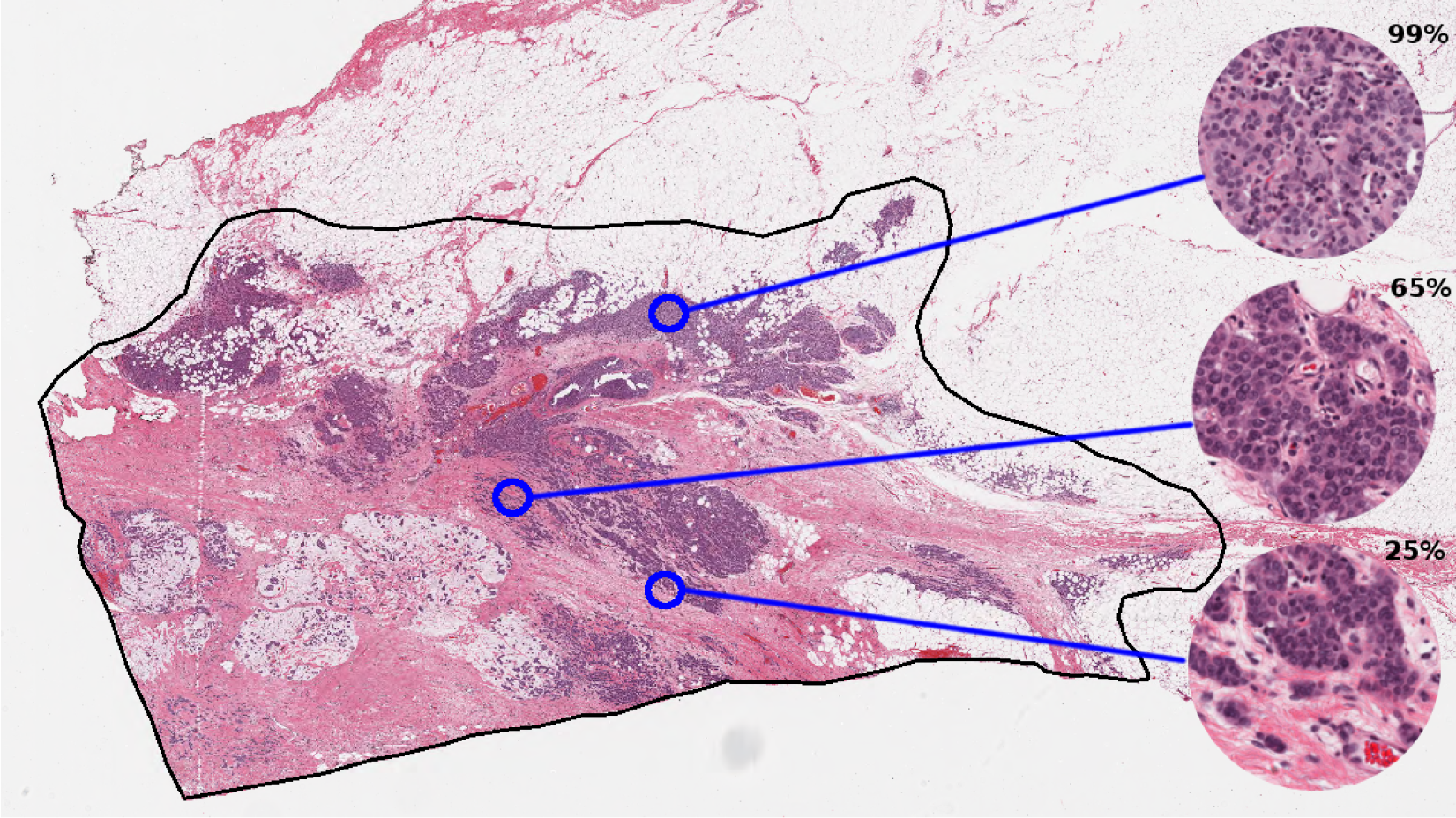
TB outlined in black in a digital slide scanned at 20X magnification (displayed at lower resolution). Regions of interest are shown in a higher magnification on the right alongside TC scores provided by an expert pathologist.

**Figure 2:**
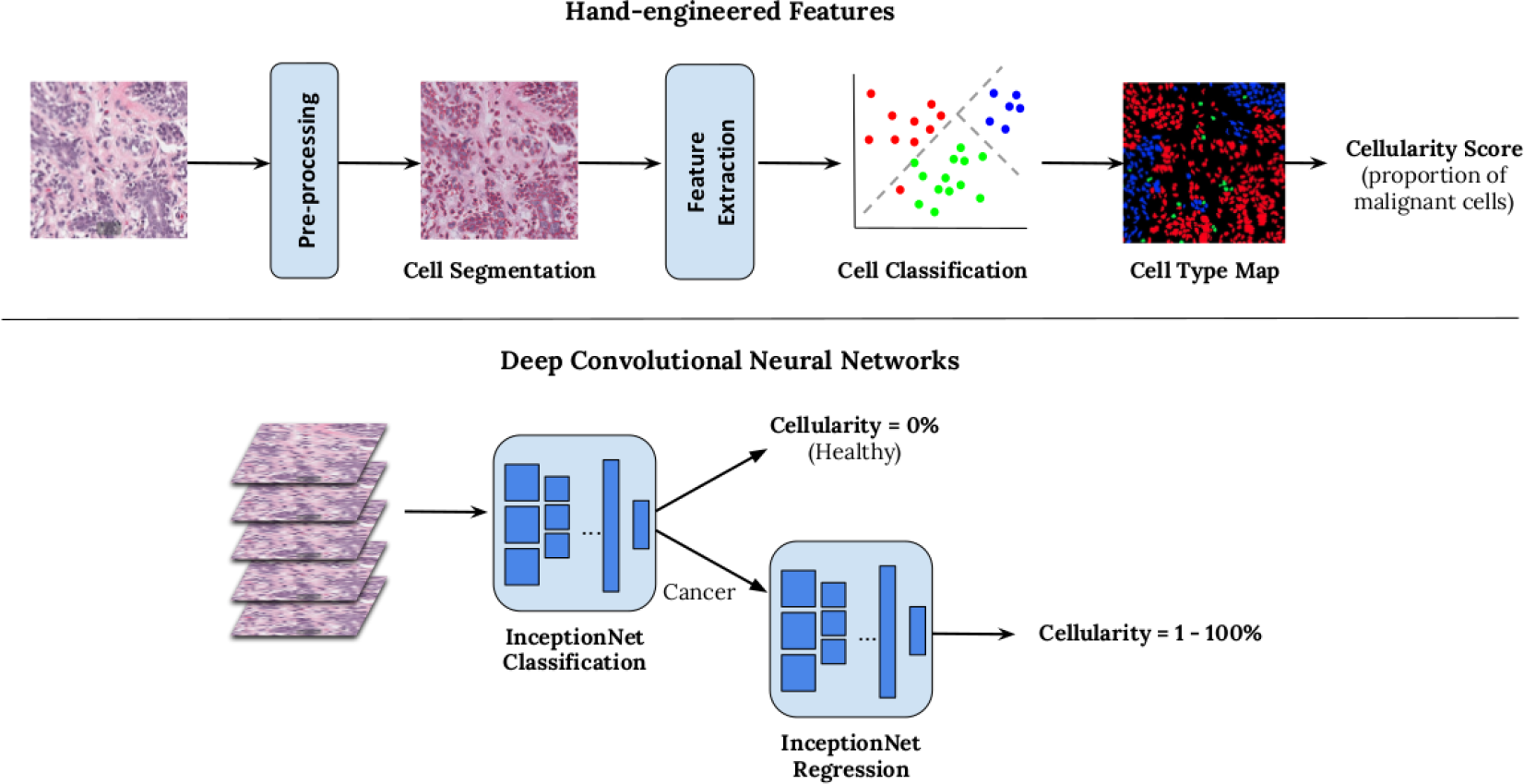
Overview of two methods for generating automated TC score. Hand-engineered feature approach is shown above and a cascade approach using two deep convolutional neural networks below.

We performed detailed comparisons between the above approaches to validate the feasibility of automation in the pathology workflow and measure the progression of image analysis techniques over the last few years to perform RCB assessment, which currently relies heavily on expert opinion. Whilst automation can also be used to locate the TB, in this paper we specifically report computation of TC with predefined TB boundaries, with the intention of implementing this as part of a larger pipeline in future work.

In this paper, we report the benefits and limitations of latest advancements in artificial intelligence, compared to a more traditional hand-engineered approach for designing algorithms. To evaluate the clinical relevance of automation on whole slide images, we also applied our trained models to high resolution digital slides scanned at 20X, achievable in a matter of minutes. We show that a localised analysis of TC on a patch-by-patch basis can be used to give a more descriptive representation of the heterogeneity of the TB area and distribution of TC scores.

## 2 Methods

### 2.1 Data

To validate different methods for computing TC, representative sections from 64 patients with residual invasive BC on resection specimens following NAT were acquired. Representative routine Hematoxylin and Eosin glass slides were scanned at 20X magnification (0.5 lm/pixel). The study was approved by the institutional ethics board. The distribution of our training and test sets were such that patient data used for training was excluded from testing. Patches, each with dimensions 512×512 pixels, were hand-selected from 96 whole slide images, 25 of which were reserved for testing purposes. In total we extracted 3,593 patches (training: 2579, test: 1121) which were then labelled manually. Patches were selected to represent a wide representation of range of TC scores.

### 2.2 Manual interpretation

An experienced pathologist with focused practice in breast pathology and a breast pathology fellow participated in the study. Each pathologist independently annotated identified patches on a digital pathology viewing platform, Sedeen Viewer (Pathcore) [8]. For each patch, a TC score, ranging from 0% to 100% for assessment of RCB [7], was provided. Patches which did not contain any tumour cells were assigned a TC score of 0%. For the test set, we retrieved two sets of annotations, from Pathologist A and Pathologist B, and compared the variability between both experts. In addition to continuous scores, both pathologists also classified each patch to low, medium, high TC and no tumour cells. Annotations were performed independently therefore each expert was unaware of scores assigned by the other.

The distribution of TC scores provided by each pathologist is given in Table 1. Note that the distribution varies considerably at higher cellularity scores (i.e. >70%) within our test sets, with 18% of Pathologist A’s scores within this range, and 31% in Pathologist B’s scores. Any automated system must be able to adjust for these differences between our experts.

**Table 1:** Number of patches in training and test set which fall into each TC score range.

|  | TC Score Range |  |  |  |
| --- | --- | --- | --- | --- |
|  | 0% | 1 - 30% | 31 - 70% | > 70% |
| Train (Pathologist A) | 701 | 840 | 665 | 373 |
| Test (Pathologist A) | 242 | 225 | 301 | 353 |
| Test (Pathologist B) | 237 | 312 | 375 | 197 |

### 2.3 Hand-engineered features

To mimic scores provided by pathologists in an automated manner, we first designed a cell nuclei segmentation algorithm to identify boundaries of individual cells of the following types: lymphocytes, epithelial cells, malignant cells. Cells boundaries were identified by removing stain variations through a series of color stretch and color space conversions. A support vector machine (SVM) classifier was trained from several appearance, textural and spatial features extracted from identified cell nuclei, producing a cell map during testing (Figure 2 (top)).

To compute a cellularity score, we aggregated the total number of cells of each cell type defined above; the proportion of malignant cells to remaining cells correlated to the automated TC score. A full description of this algorithm is given by Peikari et al. [9]

### 2.4 Deep convolutional neural network

Deep convolutional neural networks (DCNNs) are a family of architectures in artificial intelligence which are derived from a conceptual model of the human brain [10]. Typically, a DCNN consists of multiple layers, each of which contain several artificial neurons. By learning connections between hundreds or even millions of these neurons through simple linear activation functions, we can capture representations of complex data inputs. In a DCNN, groups of neurons are stacked in a series of specialized layers which can model further abstract representations of the data without manual intervention and this is where the power of DCNNs lie. Whilst there are many approaches to building DCNNs, here we opted to finetune a prebuilt network called InceptionNet [11] which has been well-adopted in the digital pathology. To compute TC scores, we trained two separate InceptionNets: one that distinguished between healthy and cancerous tissue, and the other to output regression scores on a continuous scale between 0% and 100%. Details of the implementation of the InceptionNet models is provided in supplementary material (SP1).

## 3 Results

### 3.1 Agreements between manual and automated scores

A quantitative comparison between the two trained pathologists and our two proposed image analysis pipelines are given in Table 2 in the form of intra-class correlation coefficient (ICC) values. ICC values are reported between patches extracted from digital slides and indicate variability between our readers in the independent test set. The reported intra-rater agreement between the study pathologists was 0.89. Both automated methods fell short of reported intra-rater agreements however automatically-generated scores were more consistent as shown by reported confidence intervals (shown in square brackets). This suggests that automated TC scores are consistently stable with scores retrieved from both annotators, demonstrating the advantage of reproducibility with such systems. Out of the two automated methods, DCNNs were superior, with an average agreement of 0.82, close to agreements between our experts and substantially higher than hand-engineered features. Given the upper and lower bounds of reported scores, DCNNs were on par with inter-rater agreements, and furthermore are reproducible.

**Table 2:** Two-way intra-class correlation (ICC) coefficients between two pathologists, and two automated methods for predicting TC scores. Upper and lower bounds are given in square brackets.

|  | ICC Coefficient (95% CI) |  |  |
| --- | --- | --- | --- |
|  | Pathologist A | Hand-engineered [10] | DCNN [12] |
| Pathologist A | - | 0.74 [0.70, 0.77] | 0.83 [0.79, 0.86] |
| Pathologist B | 0.89 [0.70, 0.95] | 0.75 [0.71, 0.79] | 0.81 [0.78, 0.84] |

### 3.2 Comparison between hand-engineered features and deep learning

A breakdown of prediction accuracies between both automated systems revealed that the DCNN trained to solely distinguish between health and cancerous tissue performed exceptionally well, giving accuracy rates of 93% when compared against both experts. Our hand-engineered approach fell short at 81% due to mis-identified malignant cell nuclei during the cell classification stage.

When comparing TC scores for those patches which contained cancerous structures ie TC scores >0%, we found the hand-engineered approach produced cellularity scores with strong concordance with expert pathologists (Figure 3 (left)). Whilst DCNN generated cellularity scores with good agreement with our study experts, as shown by the line of best fit, generally scores were not as precise as the hand-engineered approach. However the lack of outliers, particularly in the 0-30% range, meant that DCNN performed the best overall. The hand-engineered approach particularly suffered in the >70% range as shown in Figure 4, suggesting further work is needed to represent regions containing high proportions of tumour cells.

**Figure 3:**
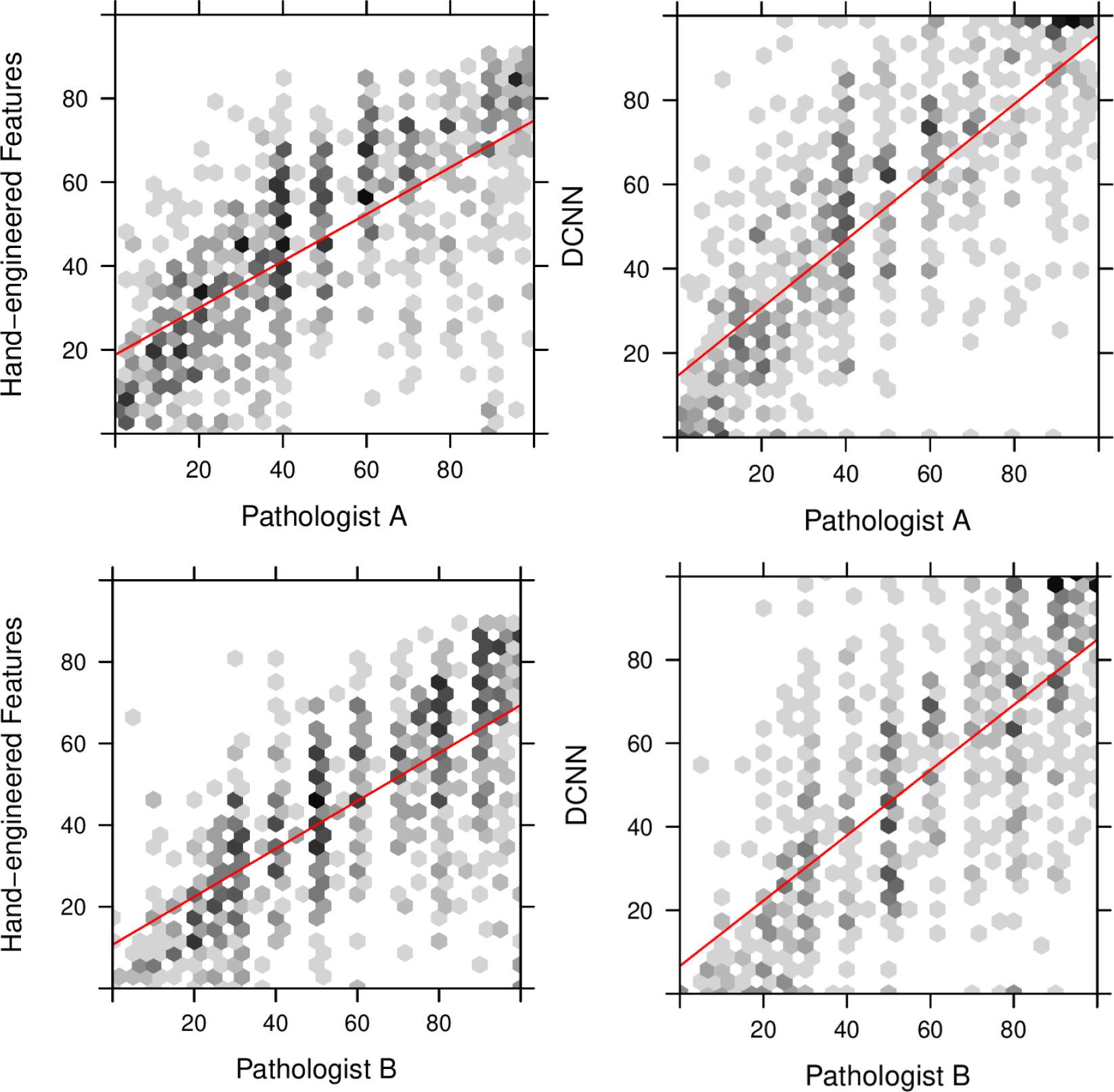
TC scores between 0% and 100% predicted by a hand-engineered approach (left) and deep neural networks (right) against scores provided by an expert pathologist (Pathologist A).

**Figure 4:**
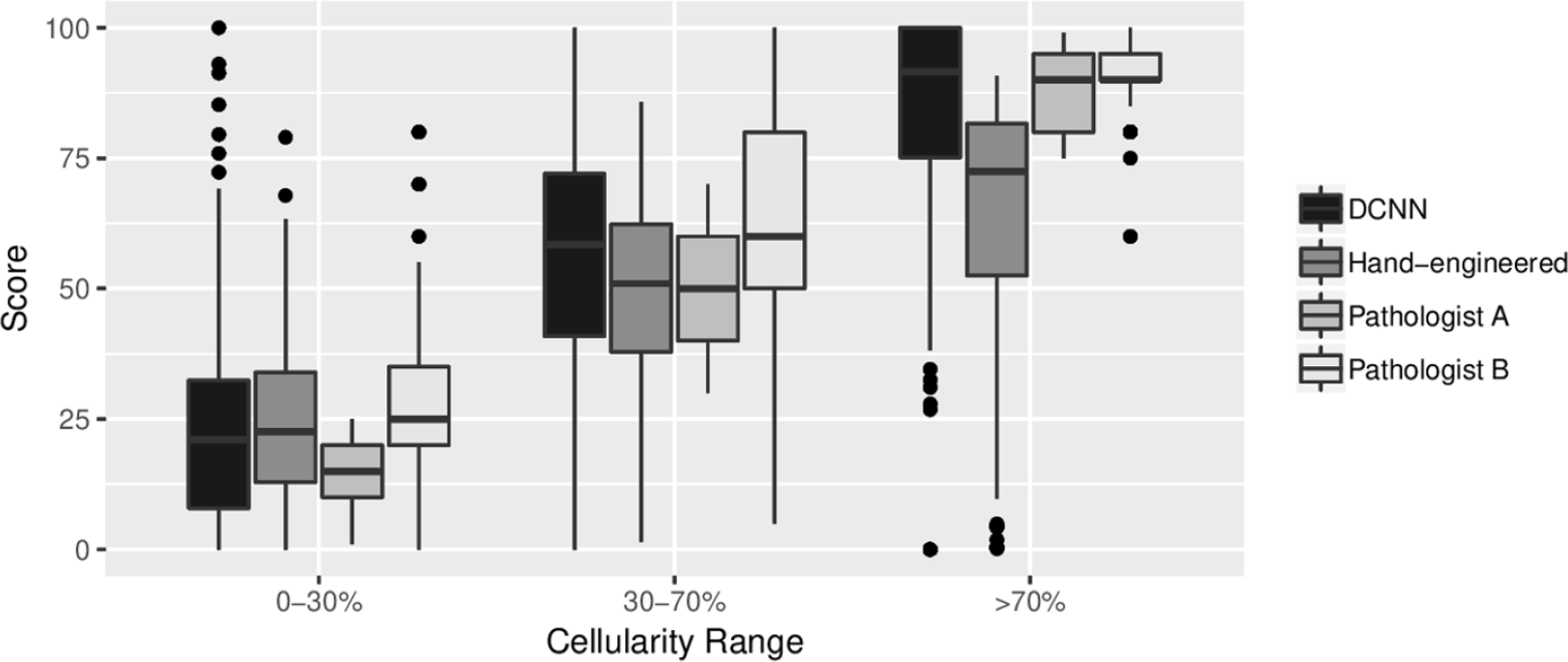
Boxplot of distribution of scores within low, medium and high ranges of TC.

## 4 Discussion

In this paper, we evaluated three methods for generating TC scores on digital slides of breast tissue for the purposes of tumour burden assessment. The standard method for computing TC scores is by visual interpretation of the TB which is a time consuming process and is limited to a rough estimate of the proportion of cancerous structures in an irregular region of interest performed by an expert. Furthermore, manual analysis is subject to inter- and intra-rater variability therefore reproducibility of TC scores is a limitation in current practice.

To increase throughput, we designed two alternative methods for generating TC scores which leverage advancements in technology to automatically analyse large whole slide images. One approach was to mimic the way in which a pathologist would compute a score, by first identifying cells in a given region of interest and then measuring the proportion of malignant to benign cells and stroma. This approach has been well adopted in the digital pathology community which has led to a large literature on cell classification and segmentation [12–14] and feature extraction methods [15].

In the last five years there has been a shift in medical image analysis to automatically extract features from image data alone using deep architectures [10]. The advantages of this approach is that there is no hand-engineering of features involved, and instead appropriate image properties are captured in a model containing several layers. There has been previous work using deep neural networks in digital pathology [16, 17], and comparisons have shown we can achieve superior performance compared to traditional feature extraction methods [18–20]. In this study, we also found that by using deep neural networks, we could achieve strong agreements with scores produced by two study pathologists; achieving ICC agreements of 0.82, approaching the intra-rater agreement of 0.89 and with tighter upper and lower bounds, suggesting more stable measurements than can be achieved manually. Our hand-engineered approach fell short at 0.75 agreement. Given these outcomes, there is potential to use automation to alleviate the burden of manually estimating TC scores which would allow assessments such as the RCB index more manageable on a routine basis.

Upon closer inspection of our results, we also found under certain conditions the use of latest automated techniques produced TC scores more similar to our experts. A subset of the scores produced by both automated systems are shown in Figure 5. The deep neural networks (D) performed better when identifying healthy tissue (top row) and patches containing almost all cancerous tissue (bottom row). The cascade approach we adopted of training a separate cancer detector, proved to be ideal for removing healthy tissue first, giving accuracy rates of 93% when identifying patches containing only healthy structures. The advantage of the hand-engineered approach comes in distinguishing between 30% to 60% TC, and this is demonstrated in Figure 3. Identifying individual cells and then measuring proportion of malignancy has a positive effect on mimicking scores produced by experts, whereas the deep neural network either over- or under-estimated in this particular range.

**Figure 5:**
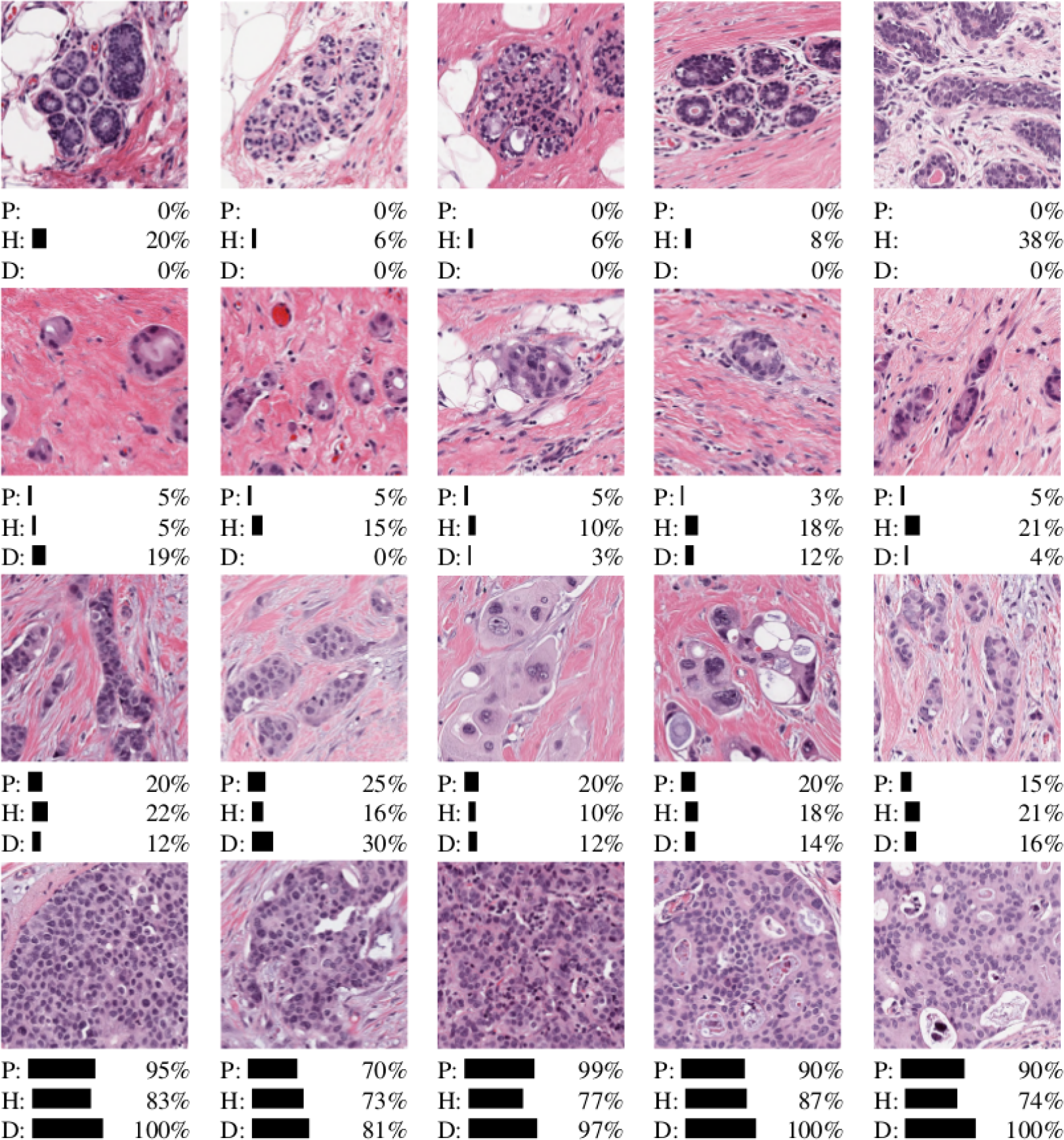
Subset of results from TC test dataset for healthy tissue, and low/medium/high TC categories (top to bottom). Scores are given for both automated systems (H = hand-engineered features, D = deep convolutional neural networks) and an expert pathologist (P).

For the purposes of determining RCB, accurate quantification of TC in the lower range leading to RCB-0 and RCB-1 classes, may enhance clinical prognostication by adopting automation. Symmans et al. [4] reported RCB-1 was a good predictor of survival outcome with 89% of triple negative breast cancer patients with RCB-1 relapse-free after 5 years; RCB-2 and RCB-3 were not prognostic. Given that the deep neural networks excelled at low TC range, the use of automation for measuring TC as a RCB component could potentially be improved by adding further training examples containing low TC scores. As it is particularly difficult to quantify TC manually, automation offers an easier method for achieving precise scores which can further contribute to use of the RCB index as opposed to RCB categorical readings.

It should be noted that whilst both automated methods reported here output scores on a continuous scale, the scores provided to the systems during training were not. Manual assessment was performed by providing an estimate of the proportion of carcinoma in each patch, often to the nearest 5% in our experiments; some variations between automation and pathologists’ scores can be explained by the scoring protocol.

Whilst here we specifically evaluated TC, the RCB index also encompasses a measure of the TB area [7]. Assessment of TB size relies upon consideration of preNAT imaging, gross examination and expert interpretation of the TB. In the current study, TC was assessed in patches derived from predefined TB regions. As such, this work is only an initial step in automating the entire RCB calculation pipeline. Further work is needed to identify “TB” and to distinguish between invasive and in situ carcinoma in a fully automated pipeline. This may require assessment of multiple digital slides per patient.

One of the main advantages of using automation is the ability to perform detailed analysis across entire whole slide images to give further contextual information. An example of our trained deep neural networks applied to whole slide images is shown in Figure 6, as heatmaps overlaid on original digital slides. We have appended a higher resolution image in supplementary materials (SP2). Blue overlays denotes low TC and red denotes patches with high TC scores. In its simplest form, this tool can be used to navigate the reader to the most interesting parts of the tissue thus eliminating around 90% of the slide. This is a desirable property in digital pathology as the substantial portion of a pathologists’ time is performed sifting through benign tissue [21] and any method for increasing throughput has significant advantage in the pathology workflow. It is important to note that the deep neural network designed to distinguish between healthy and cancerous patches suffered when applied to whole slide images compared to our patch-based test set. During training, the model was only exposed to a small subset of healthy structures i.e. fatty tissue, folding tissue, red blood cells, and were therefore unrecognizable during testing. We anticipate that with further training with more healthy patches, such errors can be avoided. In the long term, a preprocessing phase to first identify the TB region is recommended.

**Figure 6:**
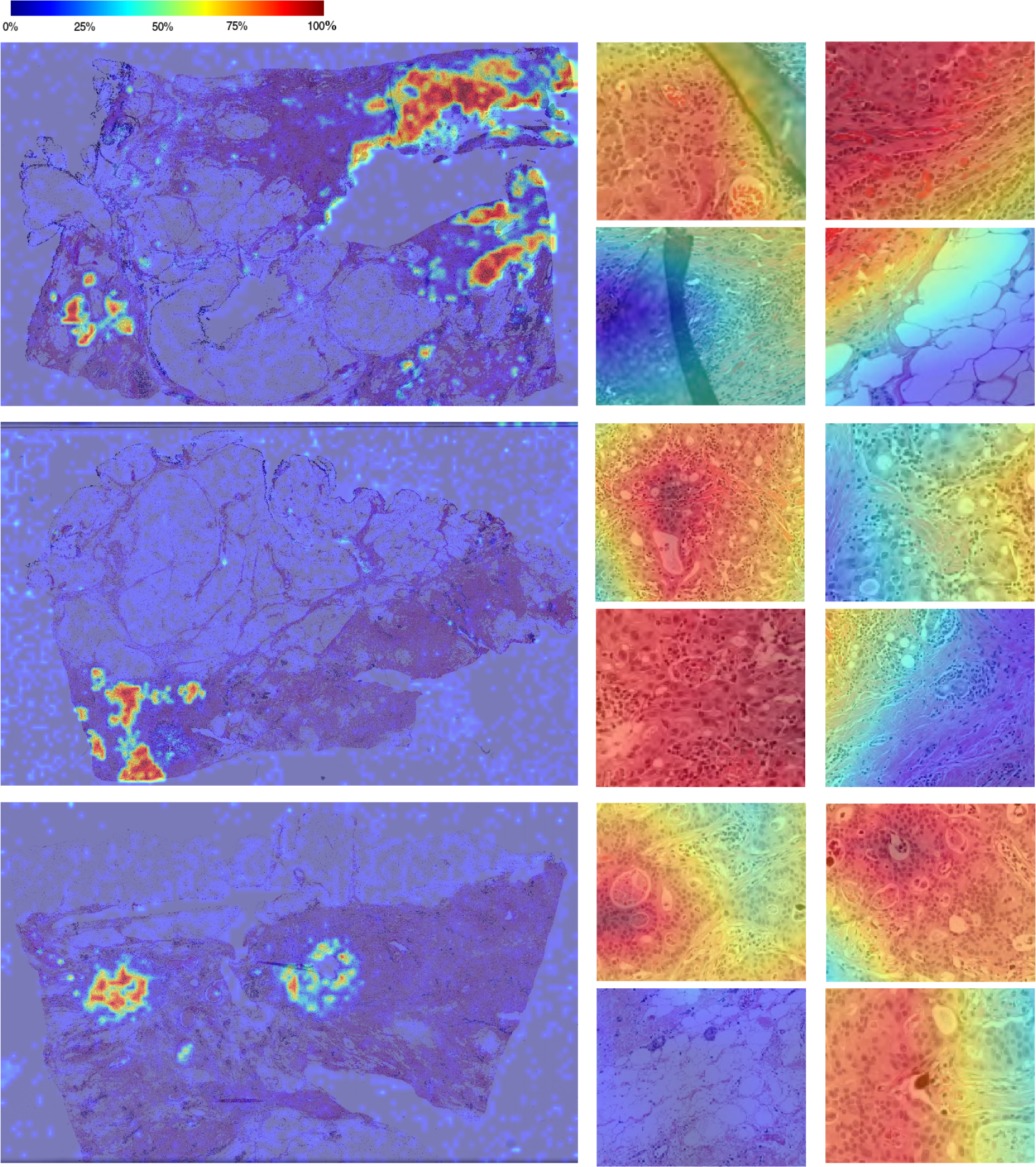
TC scores produced by a trained deep neural network overlaid on whole slide images. Scores are provided on a patch-by-patch level, where blue denotes healthy (0% TC) and red denotes 100% TC. Some close-up results of cellularity scores are provided to the right of each whole slide image.

In Figure 6, we can also see a distinct distribution of TC scores across the TB, suggesting that a global score of the entire TB may not reflect the characteristics of the TB accurately. There are clearly “pockets” of high cellularity regions and most of the TB consists of healthy or low cellularity regions. The RCB index recommends recording the average TC, however our results suggests an alternative metric which takes into account spatial distribution of TC in the TB may offer new features possibly advantageous for assessing tumour burden. Further work is also needed to investigate using continuous scores for clinical assessment, specifically the relationship between heterogeneity of the TB and prognosis.

To summarize, we performed a comparison between manual and automated assessment of TC and showed that we can gain reproducible scores with automation, and superior performance with deep neural networks. We showed that leveraging such tools on whole slide images can give us richer representations of tumour heterogeneity across the TB, and can potentially be used as an alternative metric to approximated TC scores currently used in practice.

## Acknowledgments

This work has been supported by grants from the Canadian Breast Cancer Foundation Canadian Cancer Society [grant number 703006] and the National Cancer Institute of the National Institutes of Health [grant number U24CA199374-01]. We also thank NVidia Corporation for their GPU donations for this research.

A. Martel and S. Nofech-Mozes designed and organised the research study. S. Nofech-Mozes and S. Salama annotated the dataset. A. Panah developed software to gather and store annotations for this study. S. Akbar implemented the deep learning models and M. Peikari implemented the hand-engineered image analysis pipeline. S. Akbar performed the analysis. All authors equally contributed towards the writing of the manuscript.

