## Supplementary material for "Automated and Manual Quantification of Tumour Cellularity in Digital Slides for Tumour Burden Assessment": SP1

**Supporting Information: Description of Deep Convolutional Neural Networks**

Here we provide details regarding the implementation of the deep convolutional neural networks (DCNNs) trained to generate tumor cellularity scores directly from histology image data in our study. Details regarding the setup and implementation are provided below.

To generate tumor cellularity scores, we opted to finetune a prebuilt network called InceptionNet^1^ which has been well-adopted in the digital pathology community due to the addition of “batch normalization”, which help generalizability across multiple stains and slides. InceptionNet contains a series of sub-networks of convolutional layers with various filter sizes learned in parallel which enable information to be captured at multiple scales in each stage of the network. We initialized each InceptionNet with pretrained (ImageNet) weights to stabilize the network, and then fine-tuned the last two module blocks for 30 training iterations. To closely match the dimensions of the original input data that InceptionNet was designed for, we reduced the size of each patch by a factor of two to 256 x 256 pixels. We also experimented with a windowing approach but found scaling our image performed the best (see below). We also performed a series of randomize flips, rotations and scaling to augment the dataset and increase it by 4-fold.

To compute tumour cellularity scores, we trained two separate InceptionNets: one that distinguished between healthy and cancerous tissue, and the other to output regression scores on a continuous scale between 0% and 100%. To produce these outputs from the network we performed global average pooling on the last layer of InceptionNet and added a softmax layer for classification, and a linear fully-connected layer for regression. We also added Dropout (p = 0.5)^2^ between the last and second-last layer of the network to further improve generalizability.

Our DCNN models were implemented in Keras^3^ with a TensorFlow backend and were trained on an NVidia GeForce GTX TITAN X 12GB GPU with a batch size of 32. We used the Adam optimizer to minimize training loss with a suggested learning rate of 0.001.

### **Influence of context**

To measure the influence of the field-of-view representing to the network when recognizing cancerous pathological structures, we also adapted the input to our InceptionNet model to capture less context i.e. a smaller field of view but at full resolution. We sub-windowed the original 512x512 patches into four images; keeping the number of training instances the same as in the scaled version of our input data. As tumour cellularity scores no longer reflect sub-windows of each patch, we only validated the InceptionNet which was trained to recognize cancerous structures.

The accuracy rate of the model was reduced to 89% by sub-windowing, which is a 4% reduction compared to scaling. Our results suggest capturing more contextual information through scaling is somewhat beneficial for identifying cancerous structures. This result may be due to additional image features being captured in layers of the DCNNs when a wider field-of-view is supplied. Whilst adding more training samples via augmentation helps to improve performance slightly (to 90%), this occurs regardless of the windowing scheme adapted to first extract patches from whole slide images.
