## Supplementary figures and images for "Automated and Manual Quantification of Tumour Cellularity in Digital Slides for Tumour Burden Assessment"

### SP2

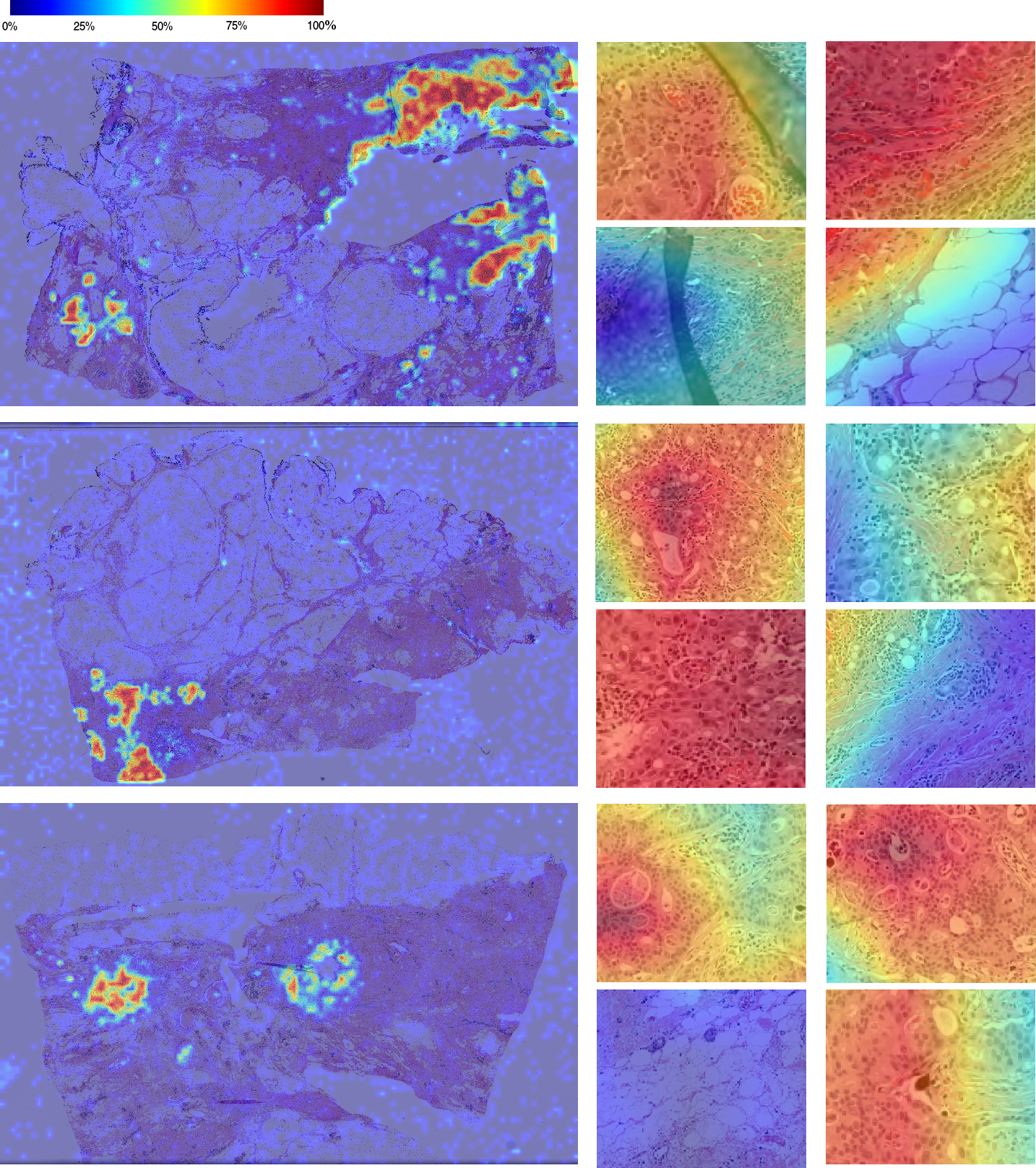
